# Structure of the integrin receptor α_M_β_2_ headpiece in complex with a function-modulating nanobody

**DOI:** 10.1101/2021.07.07.451531

**Authors:** Rasmus K. Jensen, Henrik Pedersen, Josefine Lorentzen, Nick Stub Laursen, Thomas Vorup-Jensen, Gregers Rom Andersen

## Abstract

The integrin receptor α_M_β_2_ mediates phagocytosis of complement-opsonized objects, adhesion to the extracellular matrix and trans-endothelial migration of leukocytes. Here we present the first atomic structure of the human α_M_β_2_ headpiece fragment in complex with the nanobody hCD11bNb1 determined at a resolution of 3.2 Å. The receptor headpiece adopts the closed conformation expected to have low ligand affinity. The crystal structure advocates that in the R77H α_M_ variant associated with systemic lupus erythematosus, the modified allosteric coupling between ligand coupling and integrin outside-inside signalling is due to subtle conformational effects transmitted over 40 Å. The nanobody binds to the αI domain of the α_M_ subunit in an Mg^2+^ independent manner with low nanomolar affinity. Biochemical and biophysical experiments with purified proteins argue that the nanobody acts as a competitive inhibitor through steric hindrance exerted on the thioester domain of iC3b attempting to bind the α_M_ subunit. Surprisingly, the nanobody stimulates the interaction of cell-bound α_M_β_2_ with iC3b suggesting that it represents a novel high-affinity proteinaceous α_M_β_2_ specific agonist. We propose a model based on the conformational spectrum of the receptor to reconcile these conflicting observations regarding the functional consequences of hCD11bNb1 binding to α_M_β_2_. Furthermore, our data suggest that the iC3b-α_M_β_2_ complex may be more dynamic than predicted from the crystal structure of the core complex.

## Introduction

Integrins are integral membrane proteins, which mediates cell-cell, cell-extracellular matrix, and cell-pathogen interactions. α_M_β_2_ is an integrin receptor of the β_2_ family that participates in all three types of interactions. The receptor is also known as complement receptor 3 (CR3), CD11b/CD18, and macrophage-1 antigen (Mac-1). The two most well established functions of α_M_β_2_ are those of phagocytosis of complement-opsonized cells and trans-endothelial migration (1). The two subunits α_M_ (CD11b) and β_2_ (CD18) both consist of a large N-terminal ectodomain, a single transmembrane helix and a C-terminal cytoplasmic tail. The two chains are associated through extensive non-covalent interactions (2). The ectodomain is divided into a headpiece consisting of the N-terminal domains, and the tailpiece consisting of the membrane-proximal C-terminal domains (Figure 1A).

**Figure 1:**
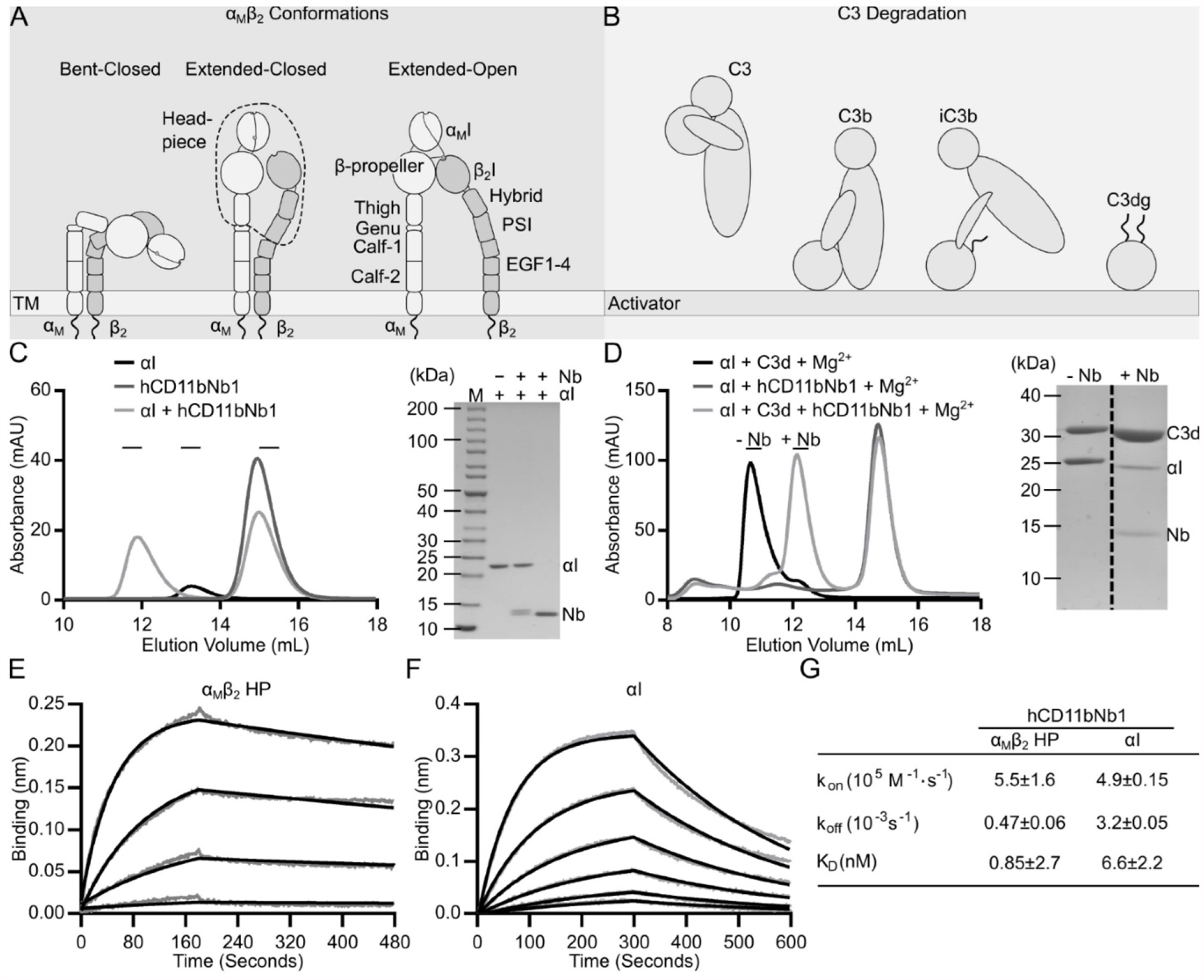
The α_M_β_2_ receptor, its iC3b ligand and characterization of hCD11bNb1. (A-B) Cartoon of the three major conformations of α_M_β_2_ and the generation of iC3b upon complement activation. The C3dg fragment encompass the iC3b thioester domain and additional disordered residues at both ends, and C3d an even smaller fragment. (C) SEC analysis of hCD11bNb1 interacting with the αI domain. In presence of the nanobody, the αI domain elutes markedly earlier. The SDS-PAGE analysis confirms the presence of both components in the early peak fraction. (D) SEC analysis demonstrates that the nanobody interferes with formation of the αI-C3d complex. (E) BLI analysis of immobilized hCD11bNb1 interacting with soluble α_M_β_2_ HP in concentrations of 50, 25, 12.5, and 6.25 nM. (F) As in panel E, but with the isolated αI domain present in concentrations of 25, 12.5, 6.25, 3.135, 1.6, and 0.8 nM. The grey curves in panels D-E are the raw data and the black curves are the fitted curves. (F) Table summarising the derived kinetic constants and their standard deviations derived from three independent experiments.

α_M_β_2_ is one of four β_2_ integrins that contain a von Willebrand factor type A domain, known as the αI domain, in their α-chain (3). The αI domain harbours a Mg^2+^ binding site, known as the metal-ion dependent adhesion site (MIDAS) (4,5). The MIDAS is directly involved in ligand recognition, where a glutamate or an aspartate of the ligand coordinates the MIDAS Mg^2+^ ion (5,6). The αI domain adopts two major conformations, open and closed (5). Transition from the closed to the open conformation leads to a rearrangement of the C-terminal α7-helix with the αI domain and a geometry of the MIDAS that allows coordination of Mg^2+^ by the ligand aspartate/glutamate (7).

During inside-out signalling, external stimuli, such as chemokines, cytokines or foreign antigens, can lead to intracellular signalling, which in turn induce conformational changes in the ectodomain that increase the ligand affinity. The ectodomain of integrins adopts at least three major conformational states with distinct affinity for the receptor ligands (3) (Figure 1A). In the low-affinity bent-closed state, the ectodomain is bent, which in the β_2_ integrins positions the αI close to the plasma membrane and the headpiece is in a closed conformation. In the intermediary extended-closed conformation, the integrin extends, leading to the αI domain pointing away from the plasma membrane, but the headpiece is still in the closed conformation. In the high affinity extended-open state, the integrin is fully extended with the β_2_ Hybrid domain swung away from the α_M_ Thigh domain.

Within the β_2_ subunit, the βI domain is structurally homologous to the αI domain, but contains two additional regulatory metal ion binding sites (8). Structure-function studies of the α_L_β_2_ (lymphocyte function-associated antigen-1, CD11a/CD18) and α_X_β_2_ (CD11c/CD18) integrins revealed that the βI domain is responsible for relaying signalling from or to the αI domain (9,10). This has led to a model for the allosteric regulation of αI domain affinity. Central to this model is that the Mg^2+^ in the βI domain MIDAS may become coordinated by Glu320 (mature numbering) from the α_M_ subunit located at the C-terminal of the αI α7 helix. Its interaction with the βI domain exerts a pull on the helix, which forces the αI domain into the open conformation during inside-out signaling. Conversely, movement of the α7 helix induced by ligand binding to the αI domain induces Glu320 coordination of the βI domain MIDAS Mg^2+^, and shifts the βI domain into the open conformation. Crystal structures indicate that the transition to the open conformation of the βI domain translates into a 60° swing out of the β_2_ hybrid domain that moves the PSI domain by 70 Å (11). This swing propagates into the open extended conformation of the β_2_ subunit, which induces intracellular signalling. Hence, signalling in integrins is bidirectional, conformational changes caused by integrin recognition of a ligand propagates through the β_2_ subunit and induces outside-in signalling (3).

One primary function attributed to α_M_β_2_ is phagocytosis of complement-opsonized cells and immune complexes. Proteolytic cleavage of component 3 (C3) by convertases deposits the opsonin C3b on the activator (Fig. 1B). The C3b has a short half-life and is quickly converted to iC3b (12) and eventually C3dg. iC3b exhibits high affinity for α_M_β_2_ whereas the minimal C3d and C3dg fragments binds the α_M_β_2_headpiece 20-fold weaker (6,13). The physiological response to iC3b recognition by α_M_β_2_ depends on both the cell type and activation state of the expressing cell, but phagocytosis of dying host cells or pathogens is the canonical response of phagocytes (14,15). α_M_β_2_ is highly expressed on the plasma membrane of myeloid cells including macrophages, monocytes, dendritic cells and neutrophil granulocytes and is upregulated from storage granules upon stimulation. Certain lymphoid leukocytes such as natural killer cells and activated T cells also express α_M_β_2_, and expression is inducible in other leukocytes (14,15). α_M_β_2_ is also highly expressed in microglia, the mononuclear phagocytes of the central nervous system (CNS), where α_M_β_2_-mediated phagocytosis of iC3b opsonized presynaptic termini of neurons is important for neural development and homeostasis (16-19). The α_M_β_2_ also plays a key role in complement stimulation of the adaptive immune system. Immune complexes containing iC3b-opsonized antigens drain into the subcapsular sinus where complement-opsonized antigens are taken up by macrophages via α_M_β_2_ and are carried across the subcapsular sinus floor (20).

Monocytes and neutrophils crawl inside the blood vessel towards a preferential site of transmigration in a α_M_β_2_ and intercellular Adhesion Molecule 1 (ICAM-1) dependent manner (21-23). Leukocytes of myeloid origin, especially monocytes and neutrophil granulocytes, uses α_M_β_2_ for diapedesis through the blood vessels to enable the cells to enter zones of inflammation. The binding to ICAM-1 presented by endothelial cells is shared with α_L_β_2_, but recent investigations show that these adhesion molecules may regulate the direction of migration in opposite ways, namely with α_M_β_2_ preventing migration against the direction of fluid flow, while α_L_β_2_, by contrast, supports such migration (24). These observations point to a complex interplay between α_M_β_2_ and α_L_β_2_, especially in the context of fluid motion exerting sheer stress on cells. Furthermore, the ability of α_M_β_2_ to bind extracellular matrix constituents such as fibrinogen and fibronectin are probably also important as step stones in the migration after leaving the blood vessel (25), and fibrinogen/fibrin as a substrate for α_M_β_2_ has long be recognized as important for leukocyte patrolling of wounds (26).

Multiple crystal structures of the ectodomain or headpiece fragments of α_L_β_2_ and α_x_β_2_ form the foundation for the mechanistic understanding of the β_2_-integrins (27-29). Studies of the α_M_β_2_ ectodomain and its complex with iC3b by negative stain electron microscopy and small angle X-ray scattering have provided low resolution structural insight about the receptor and its major ligand complex. Here we present the first atomic structure of the α_M_β_2_ headpiece fragment in complex with the nanobody (Nb) hCD11bNb1 obtained by crystallography. The receptor adopts the closed conformation with low ligand affinity. The nanobody binds to the αI domain in the α_M_ subunit in an Mg^2+^ independent manner and biophysical experiments as well as a structural comparison suggest that it acts as a competitive inhibitor of iC3b binding. In assays with cell-bound α_M_β_2,_ the nanobody in contrast, stimulates the interaction of α_M_β_2_ with iC3b. A model that considers the entire conformational spectrum of the receptor and the dynamic properties of ligand-α_M_β_2_ complexes is proposed to reconcile these conflicting observations.

## Results

### Selection and characterization of hCD11bNb1

Nanobodies are single domain antibodies of typically 120 residues derived from the variable domain of heavy chain only antibodies present in members of the camelid family. In addition to their potential for modulating the function of their antigen, nanobodies can often facilitate structure determination for challenging targets (30). Hence, we hypothesized that nanobodies specific for α_M_β_2_ headpiece could enable its structure determination and at the same time lead to identification of novel modulators of the α_M_β_2_-ligand interactions. We therefore selected nanobodies against recombinant human αI domain using a phage library generated from a llama immunized with the α_M_β_2_ headpiece. To favour the selection of nanobodies with potential for interfering with the interaction between αI domain and C3d, we performed competitive elution with recombinant C3d. This elution strategy yielded the nanobody hCD11bNb1, which we cloned into a bacterial expression vector, expressed and purified.

We validated the interaction of hCD11bNb1 with the αI domain by size exclusion chromatography (SEC) in the absence of Mg^2+^. The complex eluted markedly earlier than the two separate components and SDS-PAGE analysis confirmed the presence of both components in the early peak fractions (Fig 1C).

Next, we investigated the effect of hCD11bNb1 on the interaction between the αI domain and C3d in the presence of Mg^2+^. After having confirmed that the αI domain and C3d forms a complex eluting at 10.6 ml that is stable during SEC analysis (Fig 1D), we tested the effect of hCD11bNb1 on the interaction. In the presence of the nanobody, the peak at 10.6 ml for the αI-C3d complex disappeared. SDS-PAGE analysis confirmed that the major peak now eluting at 12 ml almost exclusively contained C3d (Fig 1D).

We next measured the binding kinetics for the hCD11bNb1-α_M_β_2_ interaction using bio-layer interferometry (BLI). We loaded His-tagged hCD11bNb1 on sensors coated with a His_5_ specific antibody and transferred the sensors into a solution containing either recombinant α_M_β_2_ headpiece (HP) or the αI domain (Fig 1E-G). Data analysis with a 1:1 binding model revealed that the nanobody binds with a low nanomolar affinity to both the α_M_β_2_ HP and the αI domain (Fig 1G). Overall, our SEC and biophysical experiments demonstrated that hCD11bNb1 bind with nanomolar affinity to the αI domain and compete with the C3d ligand, the latter a prior expectation considering the selection strategy.

### Structure determination of the α_M_β_2_ complex with hCD11bNb1

Despite extensive screening of the α_M_β_2_ HP alone and its complex with C3d or iC3b, we failed to obtain useful crystals or non-aggregated particles on cryo-EM grids. However, when bound to the nanobody, the α_M_β_2_ HP readily crystallized in a number of different organic salt capable of chelating Mg^2+^ ions. We obtained seven X-ray diffraction data sets with synchrotron radiation that extended to a maximum resolution of 3.2 Å, but all of these suffered from strong anisotropy. From any of these data sets, we were able to determine the structure by molecular replacement using the coordinates of the β-propeller from α_X_β_2_ and βI domain of α_L_β_2_ as search models. The resulting electron density and comparison with the structures of α_X_β_2_ and α_L_β_2_ allowed us to place the α_M_ Thigh domain and the β_2_ domains Hybrid, PSI and I-EGF1. Furthermore, we identified electron density that could be manually fitted with a model of the nanobody with an epitope at the αI domain. The electron density calculated from the resulting model and non-corrected diffraction data was of low quality in the receptor proximal part of the Thigh domain, the PSI and I-EGF1 and at C-terminal pole of the nanobody no matter what data set we used. Data were therefore scaled anisotropically using the STARANISO server (31), which led to a significant improvement of both 2mF_o_-F_c_ and density modified electron density maps. The data set exhibiting the best statistics after refinement of an initial model was selected for completion of the structure. To support the modelling of the α_M_β_2_-nanobody complex, we also determined the structure of hCD11bNb1 itself based on diffraction data extending to a resolution of 1.14 Å (Table 1 and Sup Fig 1). Using the anisotropy corrected data, we refined the complex structure to an R_free_ value of 0.295 (Table 1). Figure 2A-B displays the resulting structure and an example of the electron density presenting the Ca^2+^ sites in the α_M_ β-propeller. Due to our ability to compare with known structures of α_X_β_2_ and α_L_β_2_ and our high resolution structure of the nanobody, larger errors are probably not present in the model. The slightly elevated R_free_ value is therefore likely to be caused by the data anisotropy.

**Table 1.**
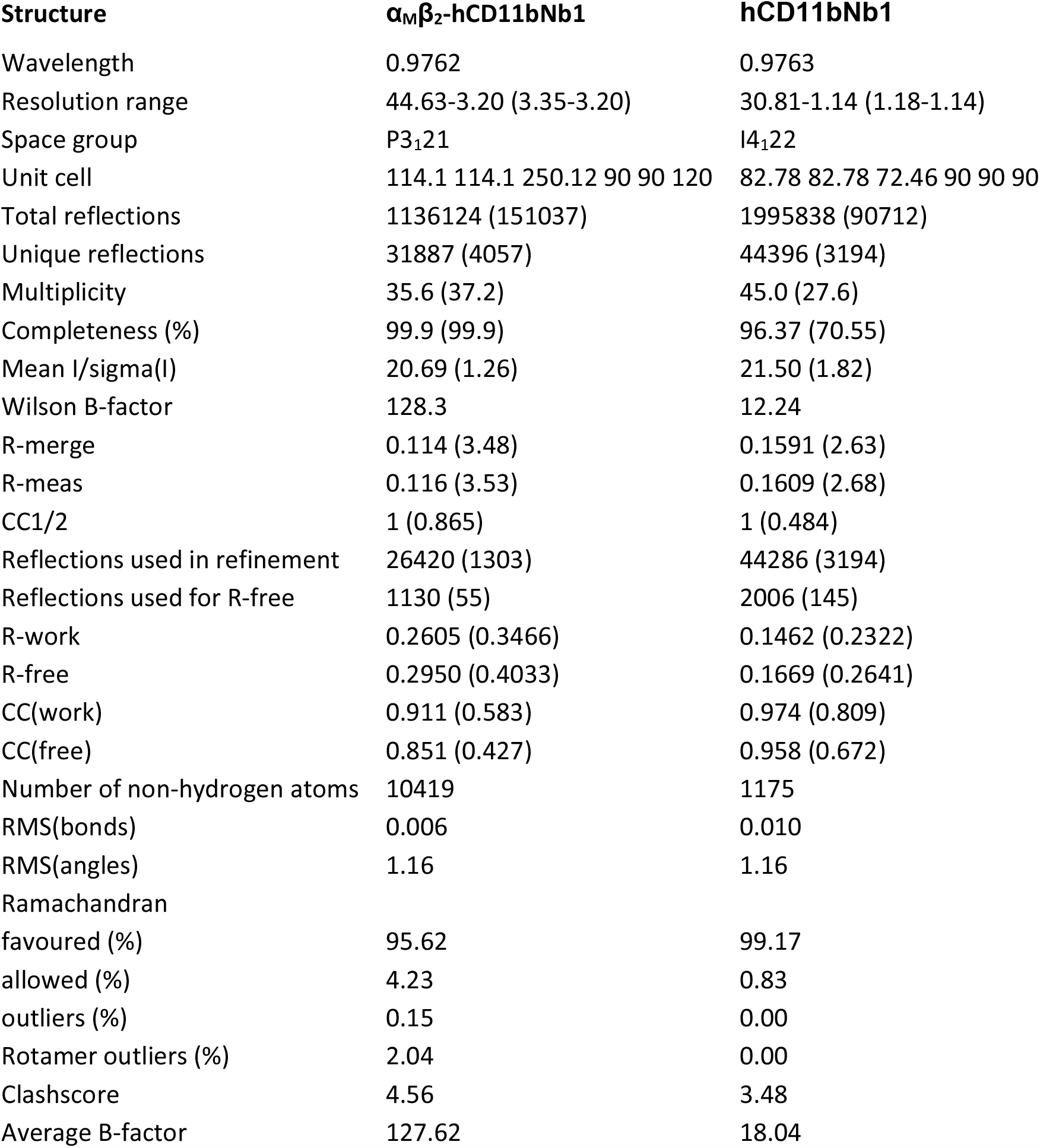
Data collection and refinement statistics. Statistics for the highest-resolution shell are shown in parentheses. For the αMβ2-hCD11bNb1 complex, the data collection statistics were calculated by XSCALE from the diffraction data not corrected for anisotropy. Refinement statistics were calculated from the data corrected for anisotropy by the STARANISO server.

**Figure 2.**
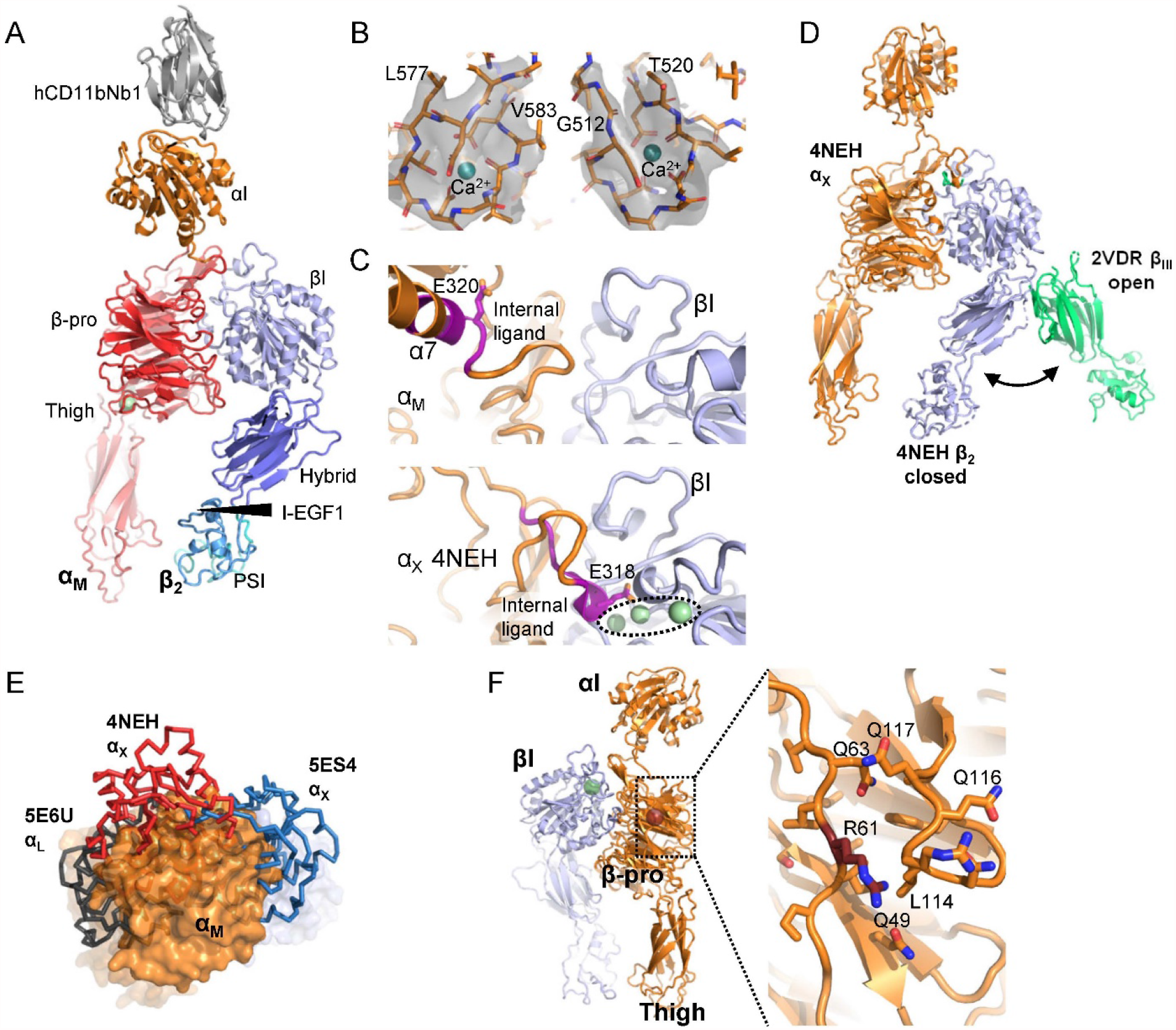
The crystal structure of the α_M_β_2_ headpiece in complex with the hCD11bNb1. (A) Cartoon representation of the structure with α_M_ domains in orange and red, βI domains in blue and the nanobody coloured grey. The promixity of β_2_ domains to the α_M_ Thigh domains demonstrates that the structure represents the closed conformation of the headpiece. (B) Omit 2mF_o_-DF_c_ electron density around the Ca^2+^ sites in the α_M_ β-propeller contoured at 1.5 σ. Residues in sticks and the ions were omitted for map calculation. (C) Comparison of the internal ligand region (magenta backbone) in our α_M_β_2_ structure and an internally liganded structure of α_X_β_2_. In the lower panel, three metal ions binding are encircled; the central ion is located in the MIDAS coordinating the internal ligand glutamate. (D) For comparison with panel A, the closed form of α_X_β_2_ (blue β_2_ subunit) is displayed together with the open conformation of the headless integrin α_2_β_III_ with a green β-subunit. (E) Compared to known structures of β2-integrins, the αI domain (orange) in α_M_β_2_ is located in a unique position. (F) Left, overall location of Arg61 40 Å from the αI domain. To the right a magnified view showing the proximity of Arg61 to the 111-118 loop that may adopt a different conformation and possess altered dynamic properties in the presence of a histidine at position 61.

### Overall structure of the α_M_β_2_-hCD11bNb1 complex

The two subunits associate through an extensive intermolecular interface formed between the β-propeller in α_M_ and the βI domain in the β_2_ subunit with a buried surface area of 3650 Å^2^. Superposition of the α_M_β_2_ onto structures of α_L_β_2_ and α_X_β_2_ reveal that the β-propeller and the βI domains interact in an almost identical manner in the three β_2_ receptors (Sup Fig 2A). Furthermore, the orientation of the Thigh domain relative to the β-propeller is also remarkably similar in α_x_β_2_ and α_M_β_2_ (Sup Fig 2B). In the α_M_ β-propeller, two calcium ions organize coordinating loop regions that together with the first residue of α_M_ forms the interface with the Thigh domain (Fig 2B). In contrast, the metal ion binding sites in both the αI and βI domains are empty and the internal ligand region at the C-terminal end of the αI α7 helix is not interacting with the βI MIDAS site (Fig 2C). The overall conformation of the α_M_β_2_ HP is closed with the Hybrid, PSI and I-EGF1 domain in the β_2_ subunit located towards the α_M_ Thigh domain, in contrast to the open conformation of the β_2_-subunit known from a structure of ligand bound α_II_β_3_ integrin (Fig 2D). Within the β_2_ subunit, the arrangement of the four domains in the closed conformation of α_M_β_2_ is also highly similar to those observed for α_X_β_2_ and α_L_β_2_ (Sup fig 2C).

### Structural basis for the altered allosteric coupling in the α_M_ R77H variant

A single nucleotide polymorphism resulting in the substitution of Arg77 to histidine in the α_M_ subunit β-propeller predisposes the carrier for systemic lupus erythematosus (SLE) (32). The R77H mutation does not change the surface expression of α_M_β_2_ on neutrophils and monocytes and inside-out signaling appears not to be affected (33,34). In contrast, the mutation does interfere with outside-in signaling since it significantly decrease phagocytosis of iC3b opsonized RBCs by macrophages and cell lines expressing α_M_β_2_ (33,34). In addition, the monocytes carrying the mutation adhere less efficiently to surfaces coated with iC3b, fibrinogen, ICAM-1, ICAM-2, and DC-SIGN. Furthermore, R77H monocytes stimulated with a TLR7/8 agonist exhibit a significantly smaller decrease in cytokine secretion upon binding of iC3b opsonized RBCs compared to wt monocytes (33). Biomembrane force probe experiments revealed that the α_M_ R77H variant of the α_M_β_2_ receptor fails to respond to force with formation of the catch bonds that would normally be induced when cells adhering through α_M_β_2_ to an immobilized ligand are exposed to an external force (35).

In our structure Arg61 (the mature numbering of Arg77 after release of the propeptide) is exposed on the edge of the α_M_ β-propeller (Fig 2F). The side chain of Arg61 only appears to interact with the nearby loop Gly111-Pro118 by non-specific van der Waals interactions. The arginine side chain does not engage in specific hydrogen bonds or electrostatic interactions that could directly explain the altered allosteric coupling in α_M_β_2_ containing the R77H variant. In our crystal structure, Arg61 is located ∼40 Å from both the α7 helix in the αI domain and the MIDAS site in the βI domain. Hence, a direct interaction of Arg61 with residues in the α_M_-β_2_ interface involved in the allosteric coupling between ligand binding and transition to the extended open conformation cannot underlie the observed functional defects. Furthermore, based on the structure of the closed-bent conformation α_X_β_2_ (27,28), we also predict that Arg61 does not interact with other domains in either of the two subunits in the bent-closed conformation of α_M_β_2_.

However, a possible consequence of a histidine at position 61 could be that the neighboring 111-118 loop (Fig 2F) change conformation and dynamic properties due to perturbation of van the der Waal interactions formed by Arg61. Alternatively, a histidine side chain at position 61 may engage in hydrogen bonds with the 111-118 loop. In support of an altered conformation of the 111-118 loop, we notice that in α_X_ with a glycine residue corresponding to α_M_ Arg61, the equivalent loop adopts a different conformation and is not in contact with the region containing the glycine. An altered conformation of the 111-118 loop could propagate and influence the dynamic properties of the N-terminal linkage between the β-propeller domain and the αI domain in residues 123-129. Alternatively, such conformational changes could propagate to α_M_ residues located at the interface to the βI domain such as the loop region Thr96-Thr101. Transmission of force is crucially dependent on a stable α_M_-β_2_ interface, even a small perturbation may give rise to the observed abnormal outside-in signaling in the R77H α_M_ variant.

### The αI domain helix 7 adopts the closed conformation

Prior structures of α_X_β_2_ and α_L_β_2_ revealed that the α-subunit β-propeller and the βI domain form a platform above which the αI domain has considerable freedom to rotate in response to crystal packing and the βI coordination state of the α_M_ internal ligand region (27-29). Confirming this idea, and in contrast to the highly conserved arrangement of the remaining domains discussed above, the orientation of the αI domain in our structure of α_M_β_2_ is unique. Compared to structures of α_X_β_2_ with the α_x_ internal ligand region interacting with the βI MIDAS, the αI domain is rotated by 180° (Fig 2E). When comparing the α_M_β_2_ structure to structures of α_X_β_2_ and α_L_β_2_ where the internal ligand region is not contacting the β_2_ MIDAS, the αI domain is rotated by 125° and 42°, respectively (Fig 2E). The α_M_β_2_ specific orientation of the αI domain may well be a result of crystal packing, since the αI-hCD11bNb1 part of the complex firmly contacts three symmetry related complexes (Sup Fig 2D). Supporting this, there are no specific interaction between the N-terminal linker region (α_M_ residues 123-131) and the platform. At the C-terminal linker region (residues 321-328) only a putative hydrogen bond between Thr322 and a sugar residue from the glycan attached to Asn375 appears to be specific for the α_M_β_2_ structure.

The αI domain has no clear density for a Mg^2+^ ion although it was present during crystallization. Also, the electron density suggests that the main chain conformation of residues Asp242-Glu244 may not be fixed, and as this region differ between the open and closed form of the αI domain (5), the conformation of the MIDAS site itself cannot be defined (Fig 3A). Hence, the nanobody does not appear to depend on a particular MIDAS conformation, and SEC analysis confirms that its binding to the αI domain is Mg^2+^ independent (Fig 1C-D). This is also consistent with that the conformation of the epitope described below does not differ significantly in structures representing the open and closed state of the αI domain.

**Figure 3.**
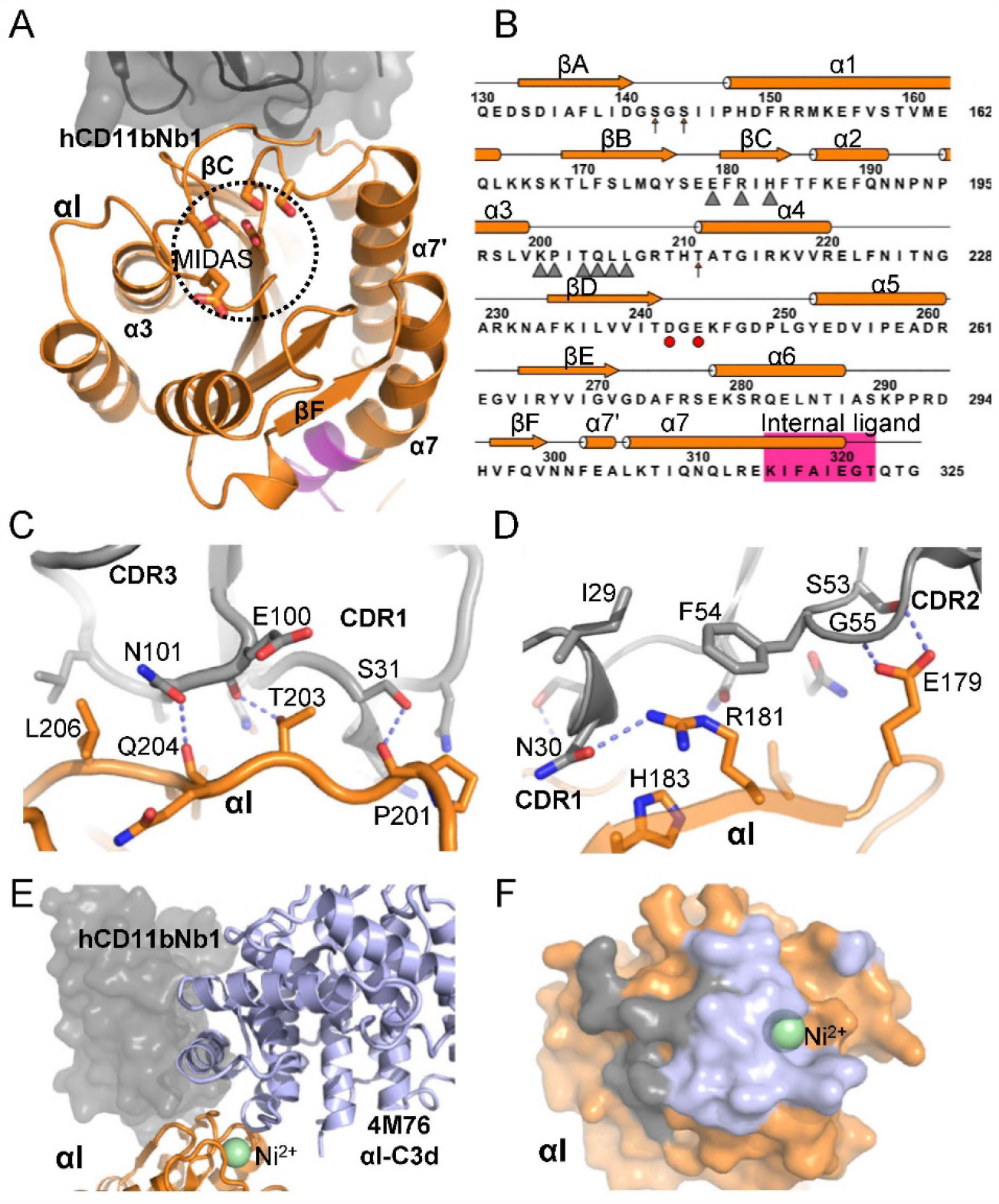
The hCD11bNb1 epitope is located adjacent to the *α*I MIDAS. (A) Top view of α_M_β_2_ αI domain with the nanobody bound with the empty MIDAS outlined. Notice also the opposite location of the nanobody relative to α-helix α7 undergoing large conformational changes during transitions between the closed and open conformations. (B) Secondary structure and sequence of the αI domain, epitope residues are marked by triangles. MIDAS residues are outlined with arrows and red circles. (C-D) Details of the interaction between the αI and the nanobody. Dashed line indicate putative hydrogen bonds. (E) Comparison of the hCD11bNb complex with our prior structure of the core complex αI-C3d created by superposition of the αI domains (orange). A small but significant overlap between C3d (blue cartoon) and hCD11bNb1 (grey surface) is predicted for a ternary complex. (F) The footprint of C3d (blue) on the αI domain is continuous with the nanobody footprint (grey) on the αI domain.

Another signature of the αI domain conformational state is the length and position of the α7 helix (5). In the nanobody complex, the α_M_ Phe302-Glu320 region is in a helical conformation (Fig 3A-B and Sup Fig 2E) meaning that this region adopts the closed conformation that prevents Glu320 from interacting with the βI MIDAS. Since hCD11bNb1 and the α7 helix are located oppositely on the αI domain (Fig 3A) and the nanobody apparently does not induce a specific conformation of the αI MIDAS, the nanobody is unlikely to influence the conformation of the α7 helix significantly. Its closed conformation is more likely to be a result of the crystal packing that favours the overall closed conformation of the β_2_ subunit in the βI domain incompatible with binding of the α_M_ internal ligand region. In summary, both the αI domain α7 helix and the overall conformation of the α_M_β_2_ HP signify the closed conformation, which appears not to be a consequence of the nanobody. In solution, the α_M_β_2_ HP contains a mixture of the open and closed conformation (13) that are likely to bind the nanobody with very similar affinities.

### The nanobody epitope on the αI domain is proximal to the C3d binding site

The quality of the electron density for the nanobody-αI interface is overall good considering the resolution and data anisotropy. Furthermore, known structures of the αI domain and our own 1.14 Å resolution structure of hCD11bNb1 itself (Sup Fig 1A-B) considerably facilitated modelling of the intermolecular interface. The buried surface area of the interface is 1330 Å^2^, which is lower but not unusual when compared to most other nanobody-antigen complexes (36). The interface is dominated by polar interactions (Fig 3C-D). Extensive burial of hydrophobic side chains at the nanobody-antigen interface plays a prominent role in our complexes of nanobodies with complement C3b, C4b and C1q (36-38) but is not observed in this case, although the hCD11bNb1 Phe54 stacks with the guanidinium group of αI Arg181 (Fig 3D). The epitope of the nanobody comprises two distinct regions in the αI domain. First, nanobody complementarity-determining regions (CDRs) 1 and 3 recognize αI residues Pro201-Leu206 with hydrogen bonds and van der Waals interactions (Fig 3C and Sup Fig 1C). Second, αI residues in the region Glu179-His183 encompassing β-strand C interact with hCD11bNb1 CDR1 and CDR2 through van der Waals interactions and hydrogen bonds (Fig 3D and Sup Fig 1C).

Interestingly, a comparison of the αI-CD11bNb1 and the αI-C3d (6) complexes revealed a small but significant overlap between the nanobody and C3d suggesting that their binding is mutually exclusive (Fig 3E-F). Specifically, the nanobody CDR3 residues 103-107 are predicted to exert steric hindrance on C3d residues Ala1214-Lys1217 in a loop region at the end of a C3d α-helix. Since the thioester domain of iC3b is expected to bind α_M_β_2_ in the same manner (6,13), this predicts that the nanobody interferes with both α_M_β_2_-iC3b and α_M_β_2_-C3d interactions. In summary, our structural analysis defined the paratope and epitope and their interactions in details and predicted that hCD11bNb1 acts as a competitive inhibitor of iC3b and C3d binding to α_M_β_2_ by exerting steric hindrance on the thioester domain of the ligand.

### Biophysical analysis of the iC3b-α_M_β_2_ complex confirms the crystal structure

To test the prediction that hCD11bNb1 acts as a competitive inhibitor for the α_M_β_2_-iC3b interaction, we took advantage of the high affinity monovalent interaction occurring between iC3b and the α_M_β_2_ HP (13). We biotinylated the free cysteine appearing in nascent C3b upon thioester cleavage using a maleimide-biotin reagent, converted the C3b to iC3b and coupled the biotinylated iC3b to a streptavidin loaded surface plasmon resonance sensor. This strategy presents iC3b in the geometry that it would have on an activator after C3b deposition and factor I degradation as outlined in Fig 1B. We next flowed recombinant α_M_β_2_ HP over the iC3b-coated sensor in the presence or absence of a 1.5-fold molar excess of hCD11bNb1. As previously reported (13), we observed a K_D_ value of 30 nM in the absence of the nanobody (Fig 4A). In the presence of hCD11bNb1, the signal decreased to 32-55 % of the signal obtained in the absence of the nanobody (Fig 4B-C) demonstrating that the nanobody compete with iC3b for binding to the α_M_β_2_ HP.

**Figure 4.**
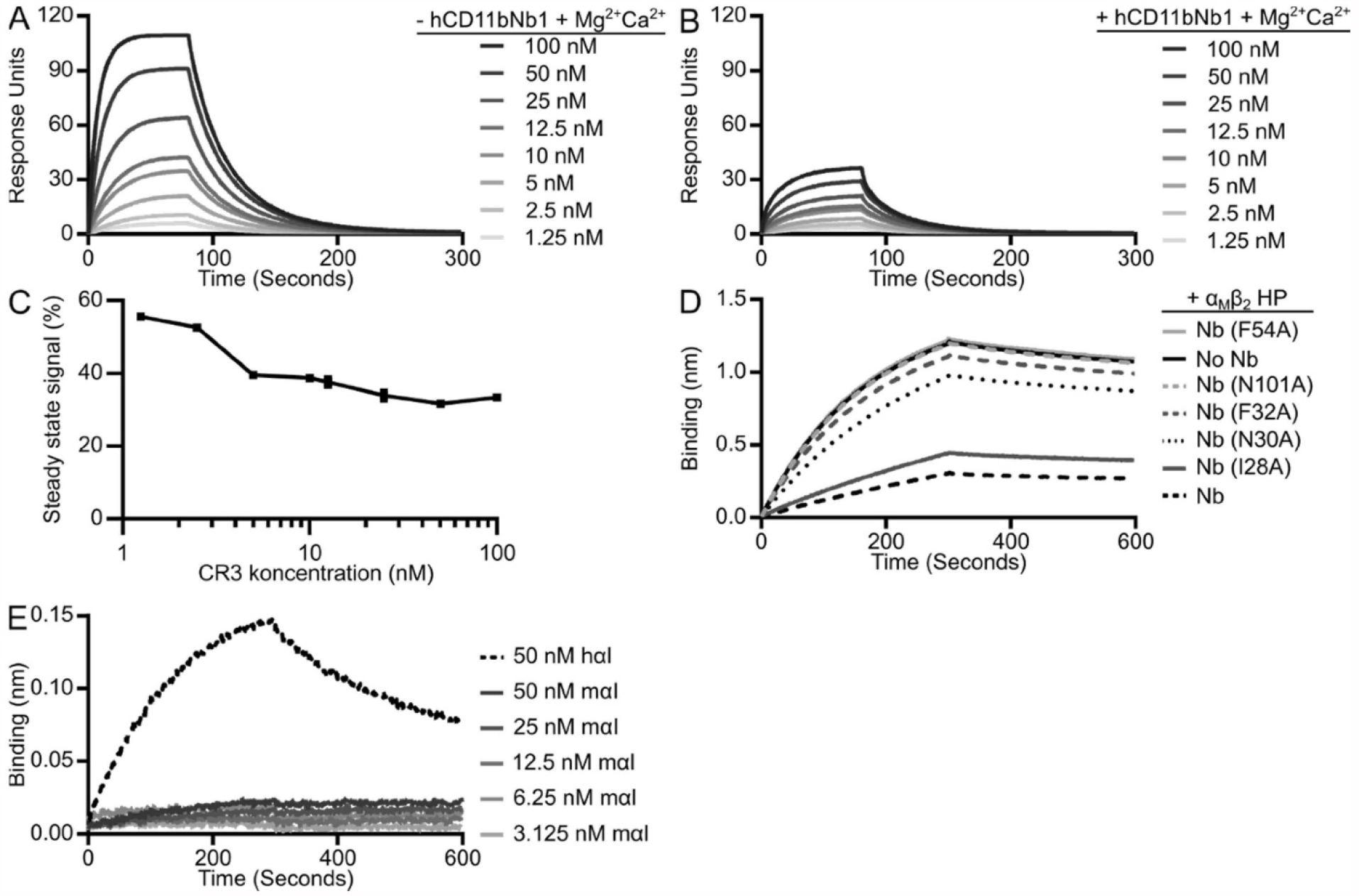
Characterization of the effect of hCD11bNb1 on the α_M_β_2_-iC3b interaction. (A-B) Biotinylated iC3b was immobilized on a streptavidin loaded SPR chip. Next, α_M_β_2_ HP in the indicated concentrations in either presence (+ hCD11bNb1) or absence (-hCD11bNb1) of a 1.5-fold molar excess of hCD11bNb1 was injected. (C) Quantitation of the decrease in α_M_β_2_ HP binding to iC3b in panel B with hCD11bNb1 present compared to panel A without hCD11bNb1 present. (D) The effect of mutations in nanobody CDRs. Biotinylated avi-tagged hCD11bNb1 was immobilized on streptavidin biosensors and the sensors were transferred into 20 nM α_M_β_2_ HP alone or 20 nM α_M_β_2_ HP pre-incubated with 10-fold molar excess of hCD11bNb1 mutants I29A, N30A, F32A, F54A, N101A or WT hCD11bNb1. (E) BLI-based analysis of the interaction between hCD11bNb1 and either human or murine αI domain. hCD11bNb1 was immobilized on anti-penta-HIS sensors and the sensors were transferred into human αI domain (hαI) at 50 nM or the murine αI domain (mαI) at 50, 25, 12.5, 6.25 or 3.125 nM.

To validate the interface between hCD11bNb1 and the α_M_β_2_ HP observed in the crystal structure, we mutated nanobody residues in direct contact with the receptor (Asn30, Phe54, Asn101) or in the vicinity (Ile29, Phe32) likely to support the conformation of the directly interacting residues. We next coupled biotinylated wild-type hCD11bNb1 to streptavidin coated BLI sensors and carried out a competition assay where variants of hCD11bNb1 were present in the fluid phase in 10-fold molar excess to the α_M_β_2_ HP. As expected, presence of the parental hCD11bNb1 in the fluid phase reduced the binding to approximately 22% of the signal obtained with the α_M_β_2_ HP only (Fig 4C). The Ile29Ala variant competed almost as well as the parental nanobody, whereas the remaining variants more or less completely lost the ability to inhibit binding of the α_M_β_2_ HP to the immobilized hCD11bNb1 (Fig 4C). Overall, our experiments with mutated nanobody variants validated the paratope-epitope interaction deduced from the crystal structure.

Since hCD11bNb1 appeared to modulate the function of α_M_β_2_, we investigated whether the nanobody could bind to the recombinant αI domain from the murine αM subunit. If hCD11bNb1 could also modulate the activity of murine α_M_β_2_, it could be an attractive reagent for *in vivo* murine models of pathogenesis where the receptor plays a role as discussed below. We charged anti-His BLI sensors with His-tagged hCD11bNb1 and compared the binding of the recombinant murine αI domain to that of the human αI domain. While we observed a strong signal for the human domain, the murine αI domain bound much weaker (Fig 4D), and the data could not be fitted to obtain rate constants and the K_D_ value. To understand why the nanobody binds the murine αI domain much weaker, we constructed a homology model of the murine domain. Inspection of this model suggested two reasons for our observations. First, a lysine in mouse αI substitutes for Thr203 in the human αI domain, this is likely to lead to steric hindrance when the nanobody binds to the murine domain. Second, L206 in the human αI domain making contact with CDR3 of the nanobody is replaced by the polar N222 in murine α_M_, which could also contribute to the decrease in affinity. In conclusion, our biophysical experiments agreed with the predictions made from the crystal structure of the complex, and demonstrated that hCD11bNb1 is unlikely to modulate the activity of murine α_M_β_2_.

### hCD11bNb1 stimulates the interaction of α_M_β_2_ presenting cells with iC3b

We next asked whether the competition between the nanobody and iC3b observed in binding experiments with isolated components and rationalized by structural comparison could be translated to the full α_M_β_2_ receptor on cells. The binding of the iC3b ligand to α_M_β_2_ was evaluated in two types of cell-based assays. The first assay was based on the binding of fluorescent-labelled iC3b (iC3b*) incubated with K562 cells that express α_M_β_2_. Here, the long incubation time permitted the reaction to reach equilibrium (13) and the fluorescent signal from cell-bound iC3b* was quantitated by flow cytometry. The signal obtained for incubations with iC3b* alone or together with hCD11bNb1 was subtracted autofluorescence (ΔMFI) measured in cells with no addition of iC3b*, either for conditions without integrin activation (Fig. 5A) or with Mn^2+^ added to activate ligand binding (Fig. 5B). In this assay, the influence of hCD11bNb1 showed a dose-dependent increase of *iC3b under conditions both without or with integrin activation (Fig. 5C-D). Importantly, an unrelated control nanobody did not increase the binding quantitatively. When this binding was normalized to the signal with no hCD11bNb1 (Fig. 5E-F), it became clear that the observed increase in iC3b* binding was independent of integrin activation: in both cases, the stimulation was approximately two-fold higher in the presence of hCD11bNb1. Especially for non-activating conditions (Fig. 5E), the binding signal followed the hCD11bNb1 concentration with lower signal for a concentration of 1 μg/ml (∼75 nM) compared to 5 or 10 μg/ml, which showed signs of saturation.

**Figure 5.**
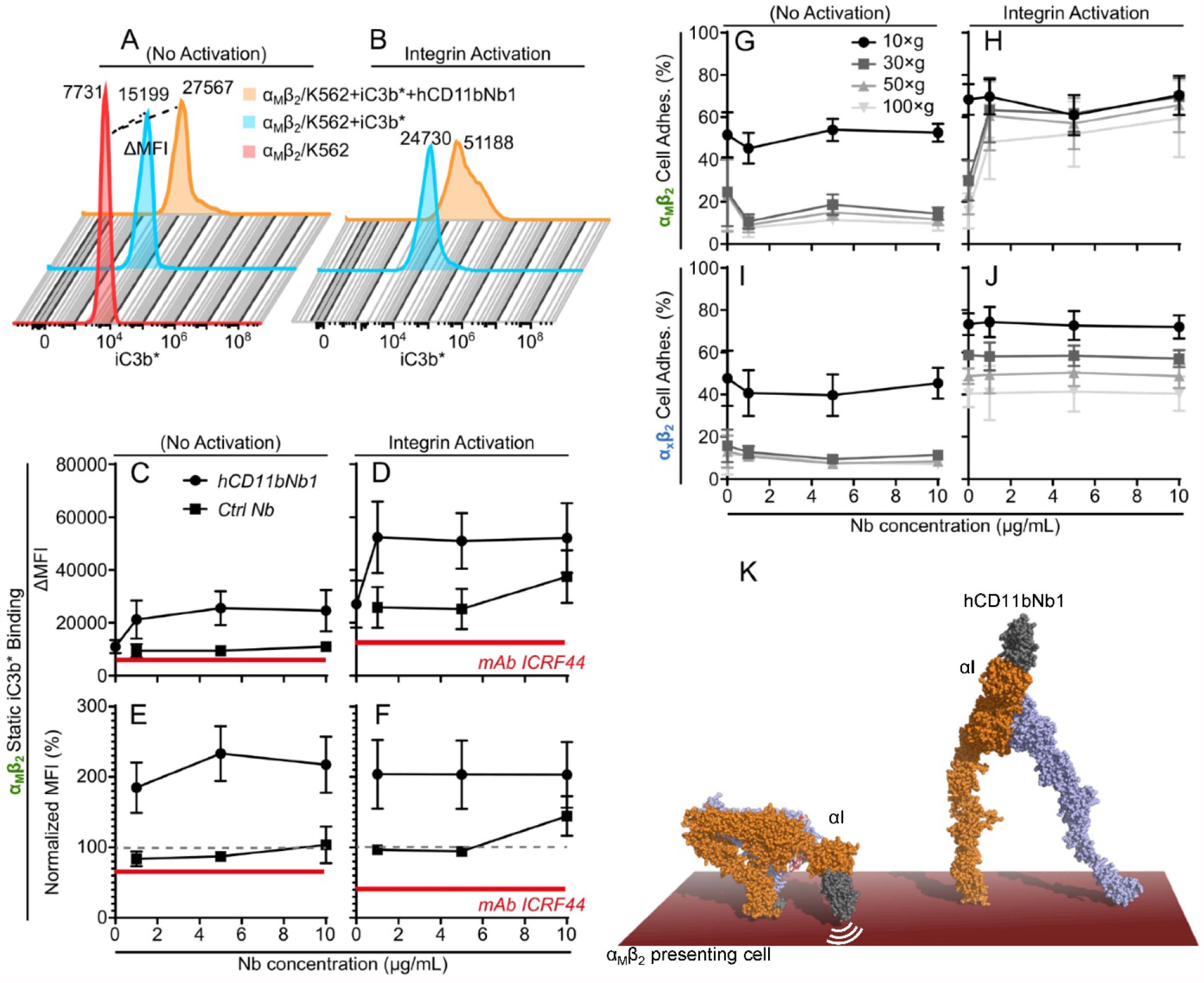
Binding of iC3b to cell-expressed α_M_β_2_ is stimulated by hCD11bNb1. The influence of hCD11bNb1 on K562 cell-expressed recombinant α_M_β_2_ ligand binding was tested in a static assay permitting equilibrium to be reached (A-F) or a forced-based cell adhesion assay (G-J), in both cases with iC3b as the ligand. (A-F) Static binding of fluorescent-labelled iC3b (iC3b*) to α_M_β_2_. (A,B) Raw data for a representative experiment with an indication of the calculation of ΔMFI by subtraction of the autofluoresence MFI (“α_M_β_2_/K562”) from the MFI for cells with iC3b* (“α_M_β_2_/K562+iC3b*”) or iC3b* and hCD11bNb1 (“α_M_β_2_/K562+iC3b*+hCD11bNb1”) under integrin resting (A) or ligand-binding activating conditions (B). (C,D) The ΔMFI was calculated for cells mixed with iC3b* and 0, 1, 5, or 10 µg/ml of hCD11nNb1, either under integrin resting (C) or activating conditions (D). As a control, non-binding nanobody was applied in the same amounts. (E,F) The data were also shown as normalized to the iC3b* without nanobody addition (≡ 100%, indicated with a grey hatched line). In Panels C-F, the influence of the function blocking mAb ICRF44 to the α_M_ chain is indicated with red, solid lines. (G-J) Influence of hCD11bNb1 on iC3b binding in a forced-based cell adhesion assay. α_M_β_2_/K562 cells were applied V-shaped wells coated with iC3b together with 0, 1, 5, or 10 µg/ml of hCD11bNb1. Increasing centrifugational force was applied in sequential steps of 10, 30, 50, and 100×*g* (indicated with grey colouring of curves). As a control, the influence of hCD11bNb1 on the binding by α_X_β_2_/K562 to iC3b was also tested as for the experiments with α_M_β_2_/K562. For each experiment in Panels C-J, the adhesion was calculated as the mean of triplicates. The results shown are the mean and SEM for three independent experiments (*N*=3). (K) Models of α_M_β_2_ on a cell membrane. The hCD11bNb1 possibly interfere with the bent closed conformation (left) due to the proximity of αI domain and the associated nanobody to the cell membrane. This may shift the conformational equilibrium of the proteins embedded in the membrane towards the extended conformation with high ligand affinity (right).

Our second assay involved force exerted through centrifugation of V-shaped microtiter wells coated with iC3b. In this way, adherent cells transmit a force load onto to the ligand-bound α_M_β_2_ mimicking physiological conditions where sheer stress acts similarly (39). In this centrifugation assay, adhesion of K562 cell expressing α_M_β_2_ to the iC3b-coated wells increased with addition of hCD11bNb1, however, with no sign of titration of the signal in the hCD11bNb1 concentration range investigated. Interestingly, in this case, the increase only occurred under integrin activating conditions in an applied force-regiment from 30-100×*g* (Fig. 5 G-H). As a control, we tested α_X_β_2_-expressing K562 cells, which also bind iC3b in the centrifugation assay. In this case, the hCD11bNb1 had no influence on the cell adhesion, consistent with the absence of the α_M_-chain (Fig. 5I-J). Overall, these cell-based assays demonstrated that hCD11bNb1, through its binding to the αI domain, can both stimulate the interaction of α_M_β_2_ with fluid phase monovalent iC3b as well as the multivalent interaction between α_M_β_2_ presenting cells and an iC3b-coated surface. Hence, these cell-based assays contradicted the model for hCD11bNb1 as an competitive inhibitor of the monovalent iC3b-α_M_β_2_ HP interaction based on our biophysical experiments and structural comparisons. In contrast, hCD11bNb1 represents a new protein based α_M_β_2_ specific agonist that promotes the interaction with iC3b presenting cells and immune complexes.

### A model for hCD11bNb1 stimulation of iC3b binding

As noted above, the hCD11bNb1 show an inhibitory effect on ligand binding to the α_M_β_2_ HP and the isolated αI domain in solution based assays. In this case, the apparent affinity (K_D_) of hCD11bNb1 was found to be in the order of 1-6 nM. By contrast, the cell-expressed α_M_β_2_ responded to hCD11bNb1 by an increase in iC3b* binding with a titration at a concentration of 75 nM, at least in the condition where no integrin activation was applied. One of the major differences between the fluid phase assays with pure components and those involving the cell-expressed α_M_β_2_ is the presence of the cell membrane in the latter experiments. Adair et al. (40) previously reported a low resolution EM structure of the α_M_β_2_ with a HP almost parallel to cell membrane, extending the bent form of the integrin sufficiently at least for smaller ligand to gain access to the α_M_ MIDAS. Furthermore, modelling of the bent conformation of α_M_β_2_ based on the structure of the α_x_β_2_ ectodomain (27,28) support that the αI domain in bent-closed α_M_β_2_ is close to the cell membrane (Fig 5K). This membrane proximity of the αI domain may contribute to the unexpected effect of the nanobody on cell-bound α_M_β_2_. Binding of the nanobody possibly promotes transition to the extended conformations (Fig 1A and 5K), and, if this is rate limiting with respect to presenting a high affinity binding site for iC3b, this alone may explain the agonist behaviour of the nanobody. This would also agree with the somewhat higher dosage requirement for inducing this effect (e.g., 75 nM) compared to the formal affinity of hCD11bNb1 for isolated HP. In the solution-based, protein interaction assays with our α_M_β_2_ headpiece, this putative membrane-dependent unbending effect on α_M_β_2_ conformation is not relevant. Here, hCD11bNb1 may either be neutral or act as a weak stabilizer of the α_M_I closed conformation, which together with steric hindrance exerted on the C3d or the iC3b thioester domain explains the observed blocking of iC3b and C3d binding. Nevertheless, if induction of α_M_β_2_ unbending accounts for the agonistic effect of hCD11bNb1, it is surprisingly that this effect is also manifest itself in the presence of Mn^2+^, which conformationally activates integrins. However, in agreement with our proposal, the influence of the Mn^2+^ is mainly through binding to a metal binding site adjacent to the βI MIDAS. Hence, occupation of this site with Mn^2+^ may not lead to a fully extended integrin, and hence, repulsion from the membrane may still be able to promote a transition to the high-affinity state of α_M_β_2_.

Despite the possible nanobody induced conformational changes in cell-bound α_M_β_2_, our comparison of crystal structures still predicts that binding of iC3b and hCD11bNb1 are mutually exclusive (Fig 3E). The discrepancy between this prediction and the cell based assay suggests that iC3b binding to the receptor is more dynamic than apparent from our crystal structure of the C3d-αI core complex (6). Considering that the overlap between the nanobody and the iC3b thioester domain only involves a few residues, it is feasible that this overlap can be reduced if the iC3b thioester domain undergo small internal conformational changes in the region involved in the overlap or the two αI binders readjust slightly relative to each other. In the light of the well-established conformational dynamics of β_2_-integrin receptors, it is plausible that the interaction of iC3b and hCD11bNb1 with the αI domain could be somewhat dynamic on the cell bound receptor compared to our crystal structures of hCD11bNb1-α_M_β_2_ and C3d-αI domain (6). Hence, complexes where the predicted overlap is reduced or eliminated may actually occur on α_M_β_2_ presenting cells. This agrees with that we and other have demonstrated that there are additional interactions outside the core complex (13) that may support a spectrum of conformations of the iC3b-α_M_β_2_ complex rather than a single rigid complex as captured in the structure of the core complex (6).

## Discussion

### The structure of α_M_β_2_ in the closed conformation

Here we present the first atomic structure of α_M_β_2_ featuring its headpiece in the closed conformation characterized by the proximity of the β_2_ Hybrid domains to the α_M_ β-propeller and the approach of the β_2_ PSI and I-EGF1 domains to the α_M_ Thigh domain. Except for the αI domain, this closed conformation of α_M_β_2_ is strikingly similar to closed conformations known from structures of α_X_β_2_ and α_L_β_2_ (27,29). Our structure of the closed conformation of α_M_β_2_ is the first step towards establishing the mechanism of outside-in signaling in this receptor. Strikingly, structures of β_2_-integrins in the open extended conformation of high ligand affinity are still lacking although low resolution negative stain EM micrographs confirm the presence of the open conformation (13,27). Comprehensive prior studies of headless integrins like α_v_β_3_ lacking an αI domain has defined in details the conformational rearrangements occurring in the βI domain upon binding of an external ligand to the βI domain. Ligand binding and modulation of the metal binding sites in the βI domain propagates into swing out of the Hybrid domain and presumably favors extension of the β_2_ subunits and its associated α-subunit and culminates in outside-in signaling (41). The β_2_ integrins including α_M_β_2_ are expected to react in a similar manner to α-subunit ligand binding and binding of the internal ligand to the βI domain.

### The αI domain in α_M_β_2_

Our comparison of the α_M_β_2_ structure with structures of α_L_β_2_ and α_X_β_2_ demonstrated a unique orientation of the αI domain relative to the platform in the closed conformation. Furthermore, two crystal structures of a bent α_X_β_2_ in which the internal ligand interacts with the βI MIDAS site demonstrated a slight variation in the orientation of the αI domain adopting the open conformation (28). The internal ligand region in these two structures is highly extended and overall these structures indicate that the distance between the MIDAS sites in the αI and βI domains as well as the orientation of the αI domain relative to the platform is not necessarily fixed (28). In crystal structures of β_2_ integrins, including our α_M_β_2_ structure, lattice packing appear to play a major role in stabilizing the position of the αI domain. Thus, it is possible that the αI domain is never locked relative to the platform in a cell bound β_2_ integrin. One striking example supporting this notion is the complex between α_x_β_2_ and its iC3b ligand, where negative-stain 2D classes revealed two opposite orientations of the ligand compared to the platform. This implies that in the ligand bound state, two orientations of the αI domain differing by up to 180° were present in the sample (42). However, other studies of α_M_β_2_ by negative stain EM featured a more defined orientation of the αI domain relative to the platform (40). Our own 3D reconstructions of the α_M_β_2_ HP also offered evidence that the αI domain is at least somewhat restricted with respect to rotation relative to the platform (13). However, because of the resolution in negative stain and the roughly spherical shape of the αI domain, it is difficult to quantitate the variability in domain orientation from such data.

Unfortunately, at present the only ligand bound structures involving αI domains are our α_M_β_2_ αI-C3d complex (6), and the complexes of α_L_β_2_ αI with ICAM-1/3/5 (7,43,44). Furthermore, a model for the α_M_β_2_ αI-GP1bαN complex based on NMR restraints and the crystal structure of the murine GPIbα N-terminal domain featured the interaction of an aspartate from a ligand α-helix (45). Detailed structures of multiple integrin-ligand complexes with intact ectodomain or their headpiece fragments are required to establish the relationship between high-affinity ligand binding, the conformational freedom of the αI domain and the structural events underlying outside-in signalling in α_M_β_2_ and other β_2_ integrin ligand complexes. To avoid crystal-packing effects on the αI location and the conformation of the rest of the receptor, single particle analysis by cryo-EM is likely to be the best approach for establishing the detailed molecular mechanism of outside-in and inside-out signalling of the β_2_-integrins.

### Function modulating molecules targeting α_M_β_2_

*In vivo* studies leave no doubt about the importance of the α_M_β_2_ as a protective agent against infection (46) and as an aggravating factor in diseases with a poorly regulated inflammatory response, for instance, as observed in animal models of multiple sclerosis and Alzheimer’s disease (47). For multiple sclerosis there is evidence from the pharmacological mode of action of drugs and animal models that α_M_β_2_ may play also in this case an aggravating role, at least in the relapsing-remitting form of the disease (48,49). With respect to stroke, blocking of α_M_β_2_ by the use of the hook worm-derived neutrophil inhibitory factor (NIF) improved the outcome in animal models (50). Later trials in humans were however compromised, by pre-existing antibodies to this parasite protein. SLE is an autoimmune disease where complement plays a central role. It is a long standing observation that α_M_β_2_ expression increases in neutrophils and scales with the severity of the disease (51). α_M_β_2_ has recently been implicated in SLE and in lupus nephritis, a kidney disease which is a common complication of SLE (52). Three missense mutations in the gene coding for α_M_ have shown a strong association with both SLE and lupus nephritis in genome-wide association studies (53-55). The negative impact on α_M_β_2_ function and strong association with SLE for the α_M_ R77H variant has been difficult to explain, but our structural data now suggest that the arginine to histidine mutation could affect the structural dynamics of the ectodomain through long-range effects on conformation and dynamic properties of residues in the α_M_-β_2_ subunit interface.

The examples of α_M_β_2_ linked diseases above demonstrate that pharmacological regulation of α_M_β_2_ activity is clinically relevant. The complications with respect to therapeutic modulation of the receptor and the repertoire of natural and man-made molecules targeting α_M_β_2_ has recently been extensively reviewed (1). Currently, the most advanced drug candidate is the α_M_β_2_ agonist Leukadherin-1 (LA-1), a small molecule that stimulates leukocyte α_M_β_2_ interaction with ICAM-1 and iC3b presenting cells (56). Mechanistically, LA-1 suppresses leukocyte infiltration into tissues by increasing α_M_/CD11b-dependent cell adhesion to ICAM-1 on the endothelium, preventing subsequent extravasation (57,58). Modelling suggests that LA-1 binds at the interface between the αI and βI domains and involves the C-terminal end of the αI α7 helix carrying the internal ligand. Such a binding pocket is difficult to reconcile with considerable rotational freedom of the αI domain in the ligand bound state, so in LA-1 bound α_M_β_2_, the αI domain may have significantly less rotational freedom compared to α_M_β_2_ not binding this small molecule drug. Our crystal structure provides a valuable scaffold for accurate modelling of α_M_β_2_ complexes with existing and future function modulating molecules.

### The mechanism and application of the hCD11bNb1 nanobody

Here we have presented the first nanobody based α_M_β_2_ agonist. With an epitope on the αI domain, hCD11bNb1 represents a highly specific reagent compared to conventional monoclonal antibodies that stimulate ligand interaction to α_M_β_2_ by manipulating the conformation of the β_2_-subunit (59). The nanobody stimulated binding of iC3b to cell bound α_M_β_2_ similar to the agonist LA-1, but with an epitope quite far from the putative LA-1 binding site at the α_M_-β_2_ interface. The mechanism of hCD11bNb1 stimulation of iC3b binding to α_M_β_2_ on cells appears to be unique and complex in the light of the inhibition of iC3b-α_M_β_2_ HP and C3d-αI interaction observed in binding experiments with the pure components.

In general, a nanobody is a versatile module that is easily humanized and targeted to specific tissues and cell types by fusion to other proteins. Fusion may also increase the short circulation time of unmodified nanobodies, reviewed in (60). Animal experiments could investigate the *in vivo* utility of properly modified hCD11bNb1 as a highly specific α_M_β_2_ agonist, but such studies are complicated by the lack of cross-reactivity with the murine αI domain. Another major complication with respect to the in vivo effects of our nanobody is the large number of proteins reported to interact with the αI domain besides iC3b (1), with ICAM-1, fibrinogen, RAGE, JAM-C, and GPIbα as prominent examples. For other ligands, steric hindrance excerpted by hCD11bNb1 could be larger than for the C3d/iC3b-αI interactions investigated here. In such cases, rather than functioning as an agonist, the nanobody may function as an antagonist. In contrast, if steric hindrance with the nanobody does not occur for other α_M_β_2_ –ligand pairs, an even stronger stimulation of ligand binding by hCD11bNb1 may be experienced.

## Experimental procedures

### Nanobody selection

The hCD11bNb1 nanobody was selected as previously described (36). Briefly, a *Lama glama* was immunized with the α_M_β_2_ headpiece by Capralogics (www.capralogics.com), and the peripheral blood lymphocytes were isolated from a blood sample. The RNA was purified from these lymphocytes, and used to prepare a cDNA library. The VHH regions were cloned into a phagemid vector by PCR, and phage display was used to select nanobodies specific towards the αI domain of α_M_β_2_. *E. coli* TG1 cells harbouring the phagemid vectors were co-infected with the VCMS13 helper phages and grown for 16 h at 30°C to generate nanobody presenting phages. Meanwhile, one well in a microtiter plate was coated with one microgram of α_M_β_2_ αI domain in 100 µL PBS, 3 mM MgCl_2_. The well of the microtiter plate was subsequently blocked by addition of PBS, 3 mM MgCl_2_ supplemented with 2% (w/w) BSA. Next, 3×10^12^ nanobody-presenting M13 phages were added to the well and the plate was incubated for 1 h at room temperature to allow binding of phages to the αI-domain. Next, the well was washed 15 times in PBS, 3 mM MgCl_2_, 0.1% Tween 20 and 15 times in PBS, 3 mM MgCl2 to remove unbound phages. The αI-domain binding phages were liberated through competitive elution by addition of a 100-fold molar excess, to α_M_β_2_ αI-domain, of recombinant C3d in PBS, 3 mM MgCl_2_. The eluted phages were amplified in the ER2748/TG1 strain E. coli and provided the basis for the second round of selection, performed similarly, however only using 0.1 µg α_M_β_2_ αI-domain for coating. ELISA was used to identify nanobodies binding the α_M_β_2_ αI-domain. To this end, an ELISA plate was coated with 100 µL of 0.1 µg/mL α_M_β_2_ αI-domain in PBS, 3 mM MgCl_2_. Meanwhile, in a 96-well format, single phage-infected colonies were inoculated LB and grown at 37°C for 6 h followed by induction of nanobody expression by addition of Isopropyl β-d-1-thiogalactopyranoside to a final concentration of 0.8 mM. The cells were grown for 16 h at 30°C, then pelleted and the nanobody-enriched supernatant was transferred to the ELISA plate. The plate was incubated for 1 h followed by 6 washed in PBS, 3 mM MgCl_2_, 0.1% tween 20. Then, 1:10.000 diluted E-tag-HRP antibody (Bethyl), was added and the plate was incubated for 1 h. The plate was washed, and developed using 3,3’,5,5’-Tetramethylbenzidine, until a clear signal was obtained. 1 M HCl was added to stop further development and the A_450nm_ was measured. Unique nanobodies were identified by sequencing and subsequently cloned into the bacterial expression vector pET-22b(+).

### Protein production

α_M_β_2_ headpiece was produced as previously described (13). In short, the supernatant of stable HEK293S cells expressing α_M_β_2_ headpiece was recovered and purified by immobilized ion-affinity chromatography using a 5 mL HisTrap Excel column (GE Healthcare). The protein was subsequently applied to a 1 mL StrepTactin column (GE Healthcare) yielding pure α_M_β_2_ headpiece. The affinity-tags and coiled coil domains, was removed by addition of the 3C protease, and a final polishing step was performed using SEC into 20 mM HEPES pH 7.5, 150 mM NaCl, 5 mM MgCl_2_, 1 mM CaCl_2_. Recombinant αI domain was purified as described (13).

C3 was purified and cleaved to C3b as described in (36). C3b was cleaved to iC3b by addition of 1 % (w/w) FH (Complement Tech) and 0.2 % (w/w) FI (Complement Tech) and the reaction was incubated for 16 hours at 4°C. The cleavage was assessed by SDS-PAGE and stopped by addition of 0.5 mM Pefabloc SC. To remove C3c or C3b, the sample was loaded on a 1 mL MonoQ column (GE Healthcare) equilibrated in 20 mM HEPES pH 7.5, 200 mM NaCl. The protein was eluted by a 30 mL linear gradient from 200 to 350 mM NaCl. C3d was purified as described (13). The hCD11bNb1 point mutants I29A, N30A, F32A, F54A, and N101A were generated by site directed mutagenesis using the QuickChange Lightning kit (Agilent). hCD11bNb1, hCD11bNb1 mutants and avi-tagged hCD11bNb1 was purified and generated as described for hC3Nb1 and avi-tagged hC3Nb1 (36). Endotoxin removal from nanobodies used for flow cytometry used for was performed as described (61). Endotoxin levels were quantified using LAL chromogenic endotoxin quantification kit (ThermoFisher) performed as described by manufacturer. Nanobodies with endotoxin levels below 2 EU/mg nanobody were considered to be endotoxin free.

### SEC assays

For analysis of hCD11bNb1: α_M_β_2_ αI interaction, 40 µg α_M_β_2_ αI domain was incubated in presence or absence of a 1.5 fold molar excess of hCD11bNb1 in 20 mM HEPES pH 7.5, 150 mM NaCl. The mix was incubated for 30 min on ice, and was then applied to a 24 mL Superdex 75 increase (GE Healthcare) column equilibrated in 20 mM HEPES pH 7.5, 150 mM NaCl. For analysis of the inhibition, 170 µg αI domain and an equimolar amount of C3d were incubated for >5 min on ice in presence or absence of a 2-fold molar excess of hCD11bNb1 in a reaction buffer containing 20 mM HEPES pH 7.5, 150 mM NaCl, 2 mM MgCl_2_. The mix was next applied to a 24 mL Superdex 75 increase (GE Healthcare) column equilibrated in 20 mM HEPES pH 7.5, 150 mM NaCl, 2 mM MgCl_2_.

### Structure determination

Crystals of hCD11bNb1 were grown by vapour diffusion at 4°C by mixing a hCD11bNb1 solution at 35 mg/mL 1:1 with reservoir solution containing 1.5 M AmSO_4_, 0.1 M BIS-TRIS pH 6.5. The crystals were soaked in reservoir solution supplemented with 30% glycerol before being flash frozen in liquid nitrogen. The data were collected at BioMAX (MAX IV, Lund) at 100 K and processed with XDS (62). A search model was prepared for molecular replacement using Phenix.sculpt (63) and the structure solved with Phaser (64). Missing residues and side-chains were built using Coot (65). In an iterative manner, the structure was rebuilt in Coot and refined with Phenix.refine using positional refinement, individual B-factors, and TLS groups. In the last round of refinement, anisotropic B-factors were refined for the sulphur atoms.

Prior to crystallization of the nanobody complex, α_M_β_2_ headpiece in 20 mM HEPES pH 7.5, 150 mM NaCl, 5 mM MgCl_2_, 1 mM CaCl_2_ was mixed with a 1.5-fold molar excess of hCD11bNb1 to a final complex concentration of 9 mg/mL. Crystals were grown at 19°C by vapour diffusion in sitting drops made by mixing the complex in a 1:1 ratio with reservoir containing 1.25 M sodium malonate, 76 mM HEPES pH 8.0, 24 mM HEPES pH 6.5, and 0.5% Jeffamine ED2001 pH 7.0. The crystals were soaked in a saturated sodium malonate solution before being flash frozen in liquid nitrogen. The data were collected at BioMAX (MAX IV, Lund) at 100 K and processed with XDS (62). The structure was determined using the coordinates of the β-propeller from α_X_β_2_ (PDB entry 4NEH) and βI domain of α_L_β_2_ (PDB entry 5E6S) in Phaser (64). The remaining domains were placed manually in Coot (65). The resulting model was refined with rigid body refinement in Phenix.refine (63). At this stage, it had become apparent that the data suffered from anisotropic diffraction. The data were therefore scaled anisotropically using the STARANISO server (31). Following this, the structure was manually rebuilt in Coot and refined with Phenix.refine using positional refinement, grouped B-factors, and TLS groups in an iterative manner. In the final round of refinement, individual B-factor refinement was conducted.

### Bio-layer interferometry

All BLI experiments were performed on an Octet Red96 (ForteBio) at 30°C and shaking at 1,000 rpm. The running and wash buffer is 20 mM HEPES pH 7.5, 150 mM NaCl, 5 mM MgCl_2_, 1 mM CaCl_2_ unless otherwise stated. For assessing the binding of hCD11bNb1 to α_M_β_2_ headpiece, anti-penta-HIS sensors (ForteBio) were first washed for 2 minutes, followed by a 5 minutes loading step where hCD11bNb1 at 5 μg/mL was loaded on the sensors. Subsequently the sensors were washed for 30 seconds, and then baselined for 2 minutes. Association to α_M_β_2_ headpiece (50, 25, 12.5, 6.25, and 0 nM) were followed for 3 minutes, followed by a 5-minute dissociation step. The binding assay with the α_M_β_2_ αI domain was performed in the same manner, except that the association was followed for 300 seconds, and that the concentrations used were 25, 12.5, 6.25, 3.135, 1.6, 0.8 and 0 nM. All experiments were performed in triplicates. The 0 nM measurements were subtracted from all data series before fitting to a 1:1 Langmuir binding model. The association was modeled as: R(t)=Rmax([α_M_β_2_]/([α_M_β_2_]+K_D_)(1-exp(-t·(k_on_·[α_M_β_2_]-k_off_))), K_D_=k_on_/k_off_, and the dissociation was modeled as a first-order exponential decay, R(t)=R(300)·exp(-k_off_(t-300 s)).

For the competition assay assessing the ability of different hCD11bNb1 mutants to compete with wild type hCD11bNb1 for α_M_β_2_ binding, the streptavidin biosensors (ForteBio) were first washed for 2 minutes in running buffer supplemented with 1 mg/mL BSA, before biotinylated avi-tagged hCD11bNb1 at 5 μg/mL were loaded on the sensors for 5 minutes. The sensors were then washed for 2 minutes in running buffer supplemented with 1 mg/mL BSA, before being baselined for 2 minutes. Thereafter, association between 20 nM α_M_β_2_ headpiece alone or 20 nM α_M_β_2_ headpiece pre-incubated with 10-fold molar excess of hCD11bNb1 mutants I29A, N30A, F32A, F54A, N101A or WT hCD11bNb1 was followed for 5 minutes. Subsequently the dissociation was followed for 5 minutes.

For analysis of the interaction between hCD11bNb1 and murine α_M_β_2_ αI, anti-penta-HIS sensors (ForteBio) were washed for 5 minutes in 20 mM HEPES pH 7.5, 150 mM NaCl. The hCD11bNb1 at 5 µg/mL was loaded onto the sensors, followed by a 2-minute wash step and a 2 minute baselined step. The association of hCD11bNb1 to murine αI (50, 25, 12.5, 6.25, 3.135 and 0 nM) or 50 nM human αI was followed for 5 minutes followed by a 5-minute dissociation step. The 0 nM measurement was subtracted from all data series. This experiment was performed in duplicates.

### Surface plasmon resonance

The SPR experiment was performed on a Biacore T200 (GE Healthcare) instrument as described (13). The system was equilibrated in running buffer 20 mM HEPES (pH 7.5), 150 mM NaCl, 5 mM MgCl_2_, 1 mM CaCl_2_. Streptavidin was immobilized on a CMD500M chip (XanTec Bioanyltics) to 200 response units. Next, biotinylated iC3b was injected on one flow cell in excess, saturating the chip surface. For competition experiment, α_M_β_2_ at 1.25-100 nM was injected in either the presence or absence of a 1.5-fold molar excess of hCD11bNb1. Upon the competition experiment, 50 mM EDTA, 1 M NaCl, 100 mM HEPES (pH 7.5) was injected over the chip to regenerate the surface.

Cell-expressed α_M_β_2_ ligand interaction with iC3b

The binding of iC3b by cell-expressed α_M_β_2_ was investigated by use of K562 cells with a recombinant expression of α_M_β_2_, or, as a control, α_X_β_2_. For binding under conditions approaching equilibrium, α_M_β_2_/K562 cells were cultured and treated with fluorescence conjugated iC3b as described (13). Briefly, the cells were kept in buffer with 1 mM Ca^2+^ and 1 mM Mg^2+^, or as further supplemented with 1 mM Mn^2+^ to activate integrin ligand binding. Following 45 min of incubation with 10 µg/ml Alexa Fluor 488-conjugated (iC3b*) together with 0, 1, 5 or 10 µg/ml of either hCD11bNb1 or an unrelated control (Ctrl) nanobody specific for complement C4 called Nb10 (66) kindly supplied by Alessandra Zarantonello. Next, the cells were briefly rinsed, followed by fixation in PBS with 0.99% (v/v) formaldehyde. As a control experiment, the well characterized function-blocking mAb to α_M_ ICRF44 (67) (Sigma-Aldrich) was also applied at 10 µg/ml. The mean fluorescence intensity (MFI) of iC3b*-bound cells was determined in a NovoCyte Flow cytometer (Agilient Technologies) by subtracting the (autofluorescence) MFI for cells with no addition of iC3b* (ΔMFI).

To investigate the influence of hCD11bNb1 under cell adhesion with a mimetic of the shear stress influencing cells under physiological conditions, adhesion of α_M_β_2_/K562 or α_X_β_2_/K562 cells were tested in a centrifugation-based assay described earlier (68). Briefly, V-shaped microtiter wells were coated with 1 µg/ml iC3b (A115, Complement Tech), or not coated as reference, and blocked in PBS with 0.05% (v/v) Tween20™. Cells were applied either in buffer with Ca^2+^ and Mg^2+^, or with a further addition of 1 mM Mn^2+^ to activate integrin ligand-binding. Nanobodies were added in concentrations of 0, 1, 5 or 10 µg/ml for either buffer condition. Following incubation for 10 min at 37°C, the cells were centrifuged at 10×*g* for 5 min and read in a fluorescence plate reader. The centrifugation and plate reading were repeated at 30×*g*, 50×*g*, and 100×*g*.

## Acknowledgements

We are grateful to excellent technical assistance from Karen Margrethe Nielsen and Bettina Grumsen and the staff at BioMAX and Petra P13 for assistance with data collection. We appreciate mentoring by Dr. Goran Bajic in the early phase of our work with α_M_β_2_

## Data availability

Coordinates and structure factor for the hCD11bNb-α_M_β_2_ and hCD11bNb1 are deposited as entries 7P2D and 7NP9.

## Author contributions

Experimental data were obtained by RKJ, HP and JL. Design of experiments and analysis of data was carried out by RKJ, TVJ and GRA. RKJ, HP, TVJ and GRA wrote the manuscript.

## Funding

This work was supported by the Lundbeck Foundation (BRAINSTRUC, grant no. R155-2015-2666), the Danish Foundation for Independent Research (grant no 4181-00137) and the Novo Nordisk Foundation Grant NNF18OC0052105.

## Supporting information

**Figure S1.**
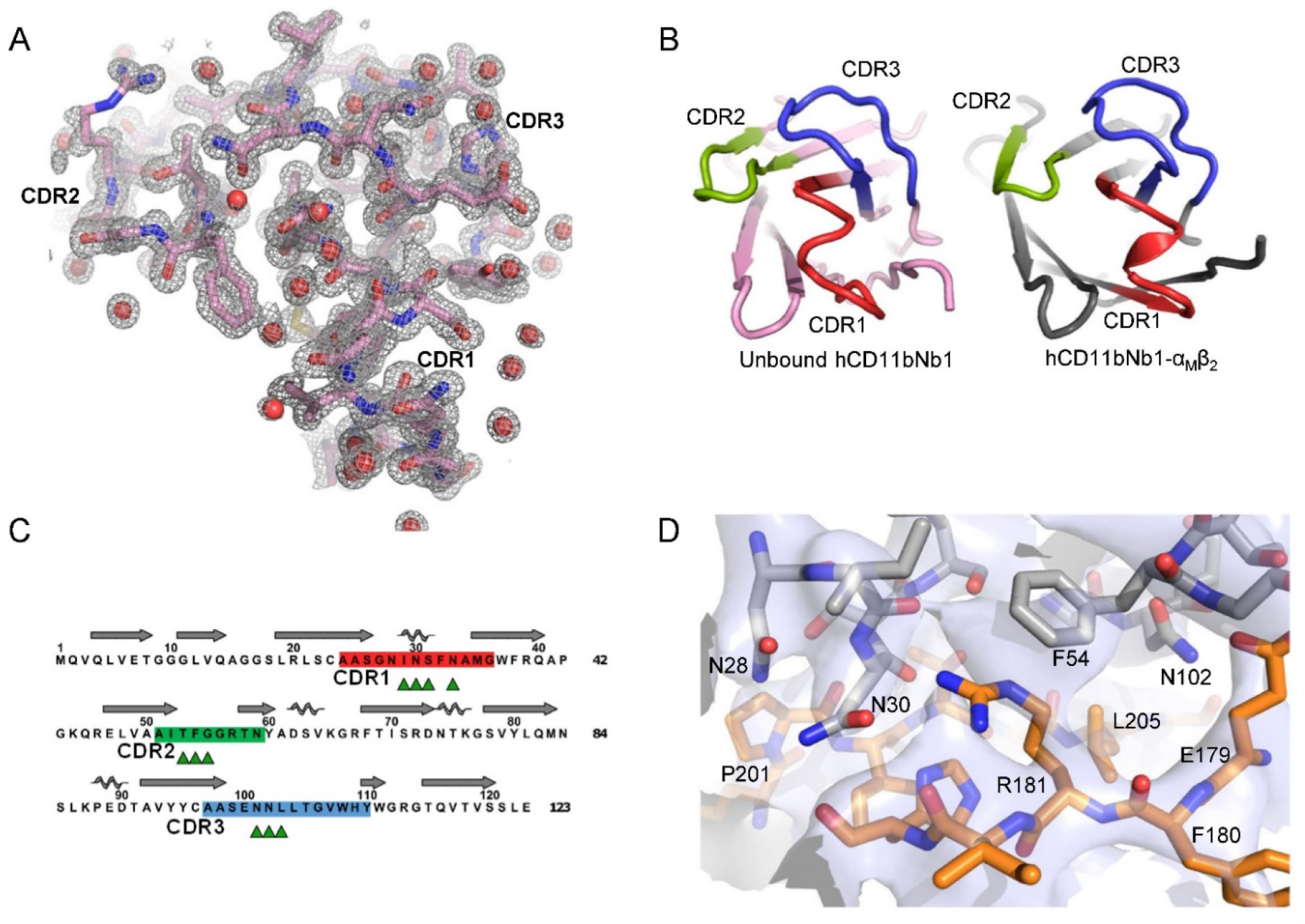
Details of the structure determination of hCD11bNb1 and its α_M_β_2_ complex. (A) Omit 2mF_o_-DF_c_ electron density around complementarity determining regions contoured at 1.5 σ for the 1.14 Å resolution structure of the unbound nanobody. (B) End view of the three CDRs demonstrating limited conformational changes upon α_M_ binding. (C) Secondary structure of hCD11bNb1 and definition of the CDRs. Triangles mark residues in the nanobody paratope contacting the α_M_ subunit. (D) Omit 2mF_o_-DF_c_ electron density for the hCD11bNb1-α_M_ interface contoured at 1.0 σ. In panels A and D, residues shown as sticks were omitted for map calculation.

**Figure S2.**
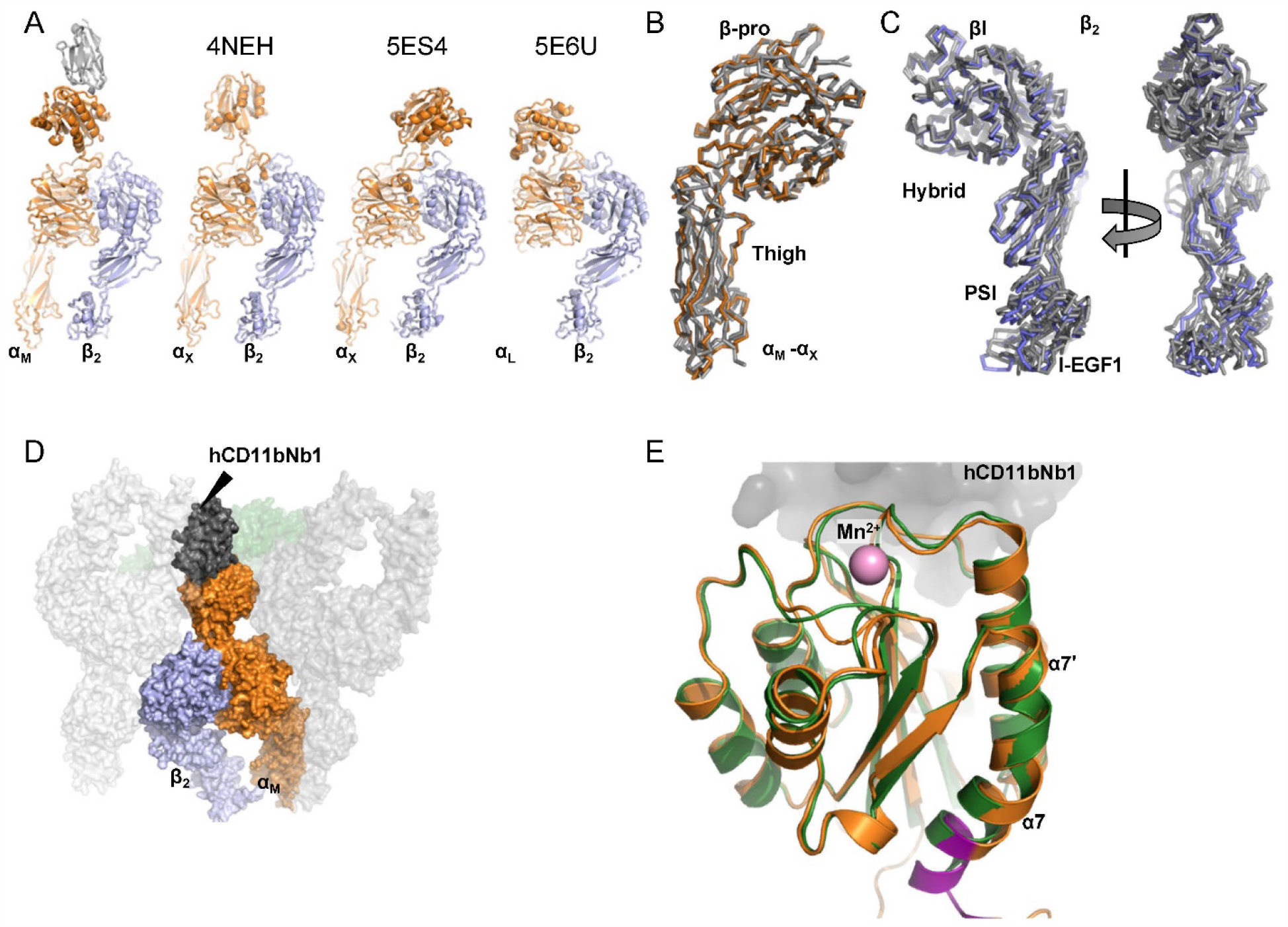
Comparison of α_M_β_2_ with known structures of β_2_-integrins. (A) A line-up of structures of the headpiece fragments (α_M_β_2_ and α_L_β_2_) and the headpiece moiety of two α_x_β_2_ ectodomain structures illustrates their strikingly similar overall conformations when aligned on their βI domains. The PDB entry is listed above each structure. (B) Alignment of α_M_ and α_x_ on their β-propeller domains demonstrates an almost identical location of their Thigh domains. (C) As in panel B with the β_2_ chains aligned on the βI and Hybrid domains for the three β_2_ integrins. Only limited variation is observed with respect to the location of the small PSI and I-EGF1 domains in the overall closed conformation of these integrins. (D) Crystal packing for α_M_β_2_-hCD11bNb1 complex showing how both the nanobody and the αI domain form contacts to three other complexes in the lattice. (E) Comparison of the hCD11Nb1-bound αI domain and the canonical closed conformation of the isolated αI domain in complex with a Mn^2+^ ion in PDB entry 1JLM.

